# Discerning amyloid network in plants

**DOI:** 10.1101/2021.01.24.427947

**Authors:** Avinash Y.Gahane, Nabodita Sinha, Talat Zahra, Ashwani K.Thakur

## Abstract

Amyloids are proteinaceous fibrillar structures and are known for their pathogenic and functional roles across the kingdoms. Besides proteinaceous deposits, amyloid-like structures are present in small metabolite assemblies and fibrillar hydrogels. Recent cryoelectron microscopy studies have shed light on the heterogeneous nature of the amyloid structures and their association with carbohydrate or lipid molecules, suggesting that amyloids are not exclusively proteinaceous. The association of amyloids with carbohydrates is further supported because the gold-standard dye of amyloid detection, Congo red, also binds to carbohydrates, probably due to similar stacking interactions. We name the association between amyloids, carbohydrates and other biomolecules as amyloid-network and propose that plants might contain such structures. Specifically, we hypothesize that cereal seeds containing glutamine-repeat-rich granules of storage proteins may have amyloid-like structures. This is because, polyQ repeats are associated with protein aggregation and amyloid formation in humans and are linked to multiple neurodegenerative conditions. Also seed storage proteins and seed cell wall proteins possess carbohydrate affinity. Thus, plant seeds might contain an intercalated network of proteins and carbohydrates, lending strength, stability and dynamics to these structures. In this paper, we show that, plant seeds have a mesh-like network that shows apple-green birefringence on staining with Congo red, a characteristic of amyloids. This congophilic network is more prominent in protein-rich seed sections of wheat and lentils, as compared to starch-rich compartments of potato. The findings suggest an amyloid network in the seeds and might be extended to other plant tissues. Further investigation with mass spectrometry and other techniques would detail the exact compositional analysis of these networks.

## Introduction

Amyloids form due to abnormal protein folding and aberrations in proteostasis. The amyloid fibrils adopt a cross-β sheet strcture irrespective of the primary amino acid sequence.[1] For clinical diagnosis, amyloid deposits are stained with Congo red dye which exhibits apple-green birefringence when visualized under a microscope equipped with polariser.[2, 3] Factors such as the protein concentration, cleavage of the protein to form amyloidogenic peptides, and the presence of amyloid enhancing factors influence the formation of these fibrils.[4] More than 30 amyloids are implicated in human and animal diseases.[5]

However, amyloids are not exclusively pathogenic[6–8] and possess structural, storage, antimicrobial, and regulatory functions.[9] Moreover, the amyloid deposits are not only made of fibrillar proteins but often contain associated proteoglycan and lipid molecules.[10–12] Alzheimer’s disease affected patient brains have exhibited lipopolysaccharides in the amyloid deposits[13], and alveolar proteinosis patient samples have shown the presence of lipid-protein aggregates in the lungs. [14] The interaction of amyloids with their surrounding biomolecules such as lipid membrane plays a role in the functional and pathological aspects of the amyloids.[15] The role of carbohydrate molecules in the deposition of amyloids has been known since more than five decades.[16] It is demonstrated that carbohydrates and especially glycosaminoglycans (GAG), play a role in promoting fibril formation due to the scaffold-like properties imparted by the GAGs.[17] Heparan sulfate, heparin, chondroitin sulfate, and hyaluronic acid are often found in amyloid deposits in the organs of AA amyloidosis patients.[18] Bacteria biofilms are also known to possess an intricate structure composed of amyloid fibres and cellulose scaffold. The resulting nanocomposite-type architecture imparts tissue-like properties such as adhesiveness and stability to the biofilm.[19] Amyloid-like structures are also formed by self-assembly of small metabolites such as phenylalanine and adenine [20, 21], suggesting that amyloid fibrilar structures are not exclusive to proteins. Also, Congo red is known to bind to carbohydrates such as cellulose and β-glucan fibrils and exhibit birefringence,[22, 23] suggesting an amyloid-like structure orientation present in these molecules.

Like the protein aggregates found in animals, plant seeds contain protein bodies formed of seed storage proteins. These are discrete, electron-rich granules that serve as nutrition storing reservoirs for seed germination.[24] Transverse sections of waxy barleys also exhibit starchy endosperm with dispersed aleurone and endosperm protein bodies in between, creating a mesh-like network.[25] A class of monocot seed proteins, prolamins composed of gliadins and glutelins are known to be rich in glutamine amino acid.[26] These proteins are deposited in the form of protein inclusion bodies similar to protein aggregates found in polyglutamine expansion disorders.[27] Wheat gluten is also known to form amyloid fibrils in certain experimental conditions.[28] There have also been reports of *in-vitro* formation of amyloid-like fibrils by seed storage proteins of maize, wheat, pea, and soybean.[29] The seed storage proteins, interestingly, are reported to possess carbohydrate-affinity due to lectin-like activity.[30] Therefore, we hypothesized that seeds might have not only amyloid fibrils but an amyloid-network composed of a scaffold of carbohydrates, lipids, and proteins. It might be possible that the presence of other biomolecules provides a framework for the seed storage proteins to accumulate and perform their functions. In this article, we have shown that the seeds contain a congophilic network, probably composed of an association of proteins and carbohydrates.

## Results and discussion

### Congo red and ThT staining of plant carbohydrates

Before studying the histology of seed sections, we prepared starch and cellulose suspensions from starch powder and microcrystalline cellulose, respectively, and the samples were stained with Congo red. Starch is a source of carbohydrates and is made of amylose and amylopectin. Short chains of amylopectin form double helices, which might crystallize and contribute to the crystallinity of the starch granules. Under polarized light, starch granules often exhibit a “Maltese cross” structure showing their radial organization.[31] Cellulose, on the other hand, is the structural carbohydrate of plant cell walls. The primary cell walls of plants are composed of long cellulose microfibrils in a matrix of glucans, pectins, and proteins. The growth of the plant cells are directional and depend on the microfibril orientation.[32] The starch suspension under bright light did not show any notable features but, under polarized light, showed blue Maltese cross-like structures. The cellulose suspension produced needle-like structures which stained red with the dye. However, no significant apple-green birefringence was observed under the cross-polarizer. This suggests that cellulose or starch granules prepared *in-vitro* do not have amyloid properties. Since Congo red is known to bind to cellulose fibrils stacked in a certain orientation in cell walls, the *in-vitro* preparations of cellulose might not show similar birefringence. However, the cellulose and starch assemblies represented here may be described as amyloid-like since these bind to thioflavin-T and emit a strong green fluorescence under fluorescent microscope.

**Figure 1:**
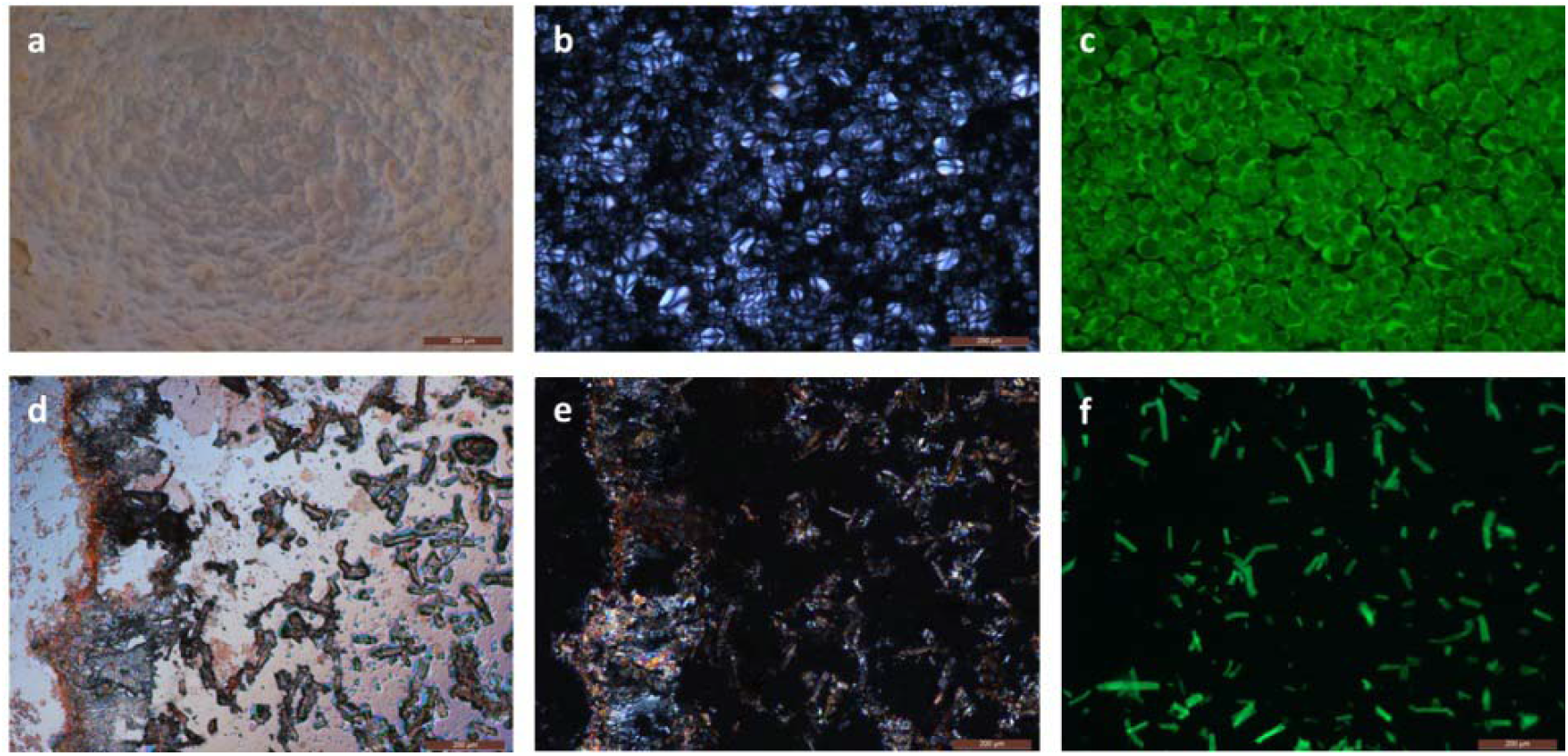
Congo red and ThT staining of starch (a-c) and cellulose (d-f). The starch suspension exhibit ‘Maltese-cross’ structures in polarized light when bound to Congo red. Starch, when stained with ThT, shows intense green fluorescence characteristic of amyloid-like structures. The cellulose suspension did not show significant apple-green birefringence on binding to Congo red. However, intense green fluorescence was observed with ThT stain. (Scale bar - 200μm)

### Congo red and ThT staining of ethanol-extracted wheat gluten

Wheat gluten consists of glutamine-rich proteins such as gliadins. We hypothesized that these glutamine-rich proteins might form protein aggregates in the seed storage vacuoles of plants. Wheat gluten is the proteinaceous mass left behind after washing wheat flour repeatedly. It is made of a mixture of proteins, belonging to the class of prolamins and is the primary storage form as protein bodies in seeds.[33] The prolamins are found to be rich in glycine, proline and glutamine amino acids. The glutamine-repeat rich nature might lead to their amyloidogenicity. In addition, wheat gluten is known to form amyloid-like structures in specific experimental conditions.[34] Ethanol extraction of gluten results in isolation of gliadins, composed of a single polypeptide chain. We prepared ethanol extract of gluten under non-reducing conditions and stained the extract with congo red to find a network-like structure, but it did not show significant birefringence under polarized light. The ThT staining of the ethanol extract showed a ribbon-like form with green fluorescence exhibiting the amyloid-like structure of gluten extract.

**Figure 2:**
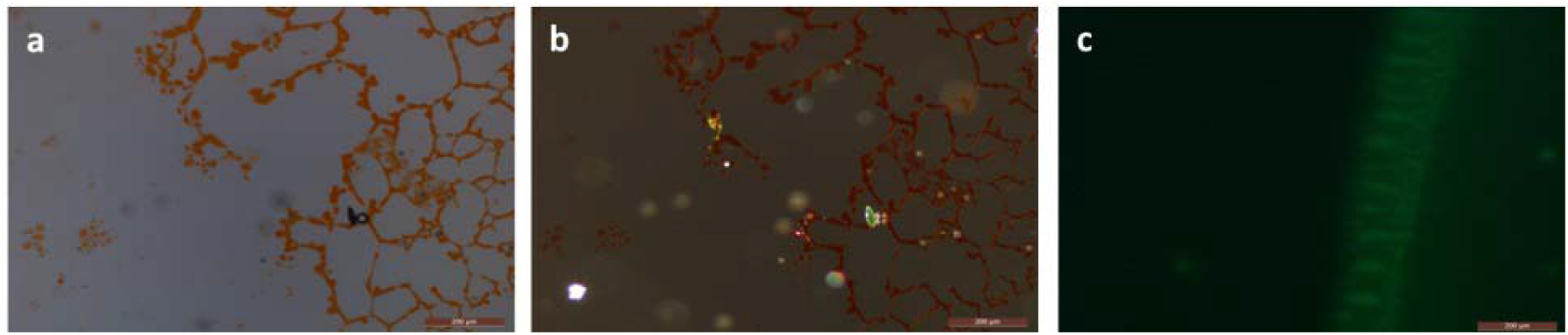
Congo red (a,b) and ThT (c) staining of ethanol extract of gluten. The gluten extract did not show any significant birefringence on staining with Congo red. ThT binding exhibits green fluorescence. (Scale bar - 200μm)

#### Congo red staining of seed sections

Congo red is a “gold standard” dye used to detect amyloids and is used in the diagnosis of amyloid deposits since their discovery. Literature suggests that seeds contain defensin proteins such as vicilin which possess amyloid properties.[35] However, Congo red staining of entire seed sections has not been performed yet to the best of our knowledge. For characterizing the Congophilic structures in the seed sections, we choose wheat (*T.aestivum*) seeds. The endosperm cell walls, nucellar projection cell walls and the seed coat showed an apple-green birefringence, characteristic of amyloids and showed amyloid-specific transformations on rotating the polarized light.

The aleurone and endosperm cell walls of wheat are composed of ~40% sugar by dry weight. Among the sugars, β-glucan and arabinoxylan are the major components while cellulose and glucomannans are present in smaller quantities.[36] The aleurone cell walls also contain hydroxycinnamic acids and ~1% protein in their dry weight. Compared to the aleurone layer, the endosperm cell walls have lesser amount of protein in them but the amino acid sequences of the proteins in both these compartments are similar. Both these types of cell walls have proteins rich in glycine, proline and serine. Literature suggests that these proteins cross-link to the polysaccharides using the tyrone-hydroxycinnamic acid dimerization and the tyrosine-tyrosine bridges present in the structures make them resistant to alkali and enzymes.[37] Another contributing factor to their stability could be that interactions of these carbohydrates and proteins lead to the formation of amyloid-like structures, resulting in their binding to the Congo red dye.

The crease region of the wheat seeds was another Congophilic region showing binding to Congo red at the cellular walls. These creases are mostly made of nucellar projections and epidermis cells through which the wheat grains imbibe water and other nutrients for germination. The cell walls of the nucellar projections are rich in arabinoxylan, mixed-linkage glucan, and homogalacturonan.[38] The nucellar cells of young embryos generally contain arabinogalactan peptides which are rich in glycosylated hydroxyproline amino acids[39] and might play a role in association with the cell wall matrix carbohydrates.

**Figure 3:**
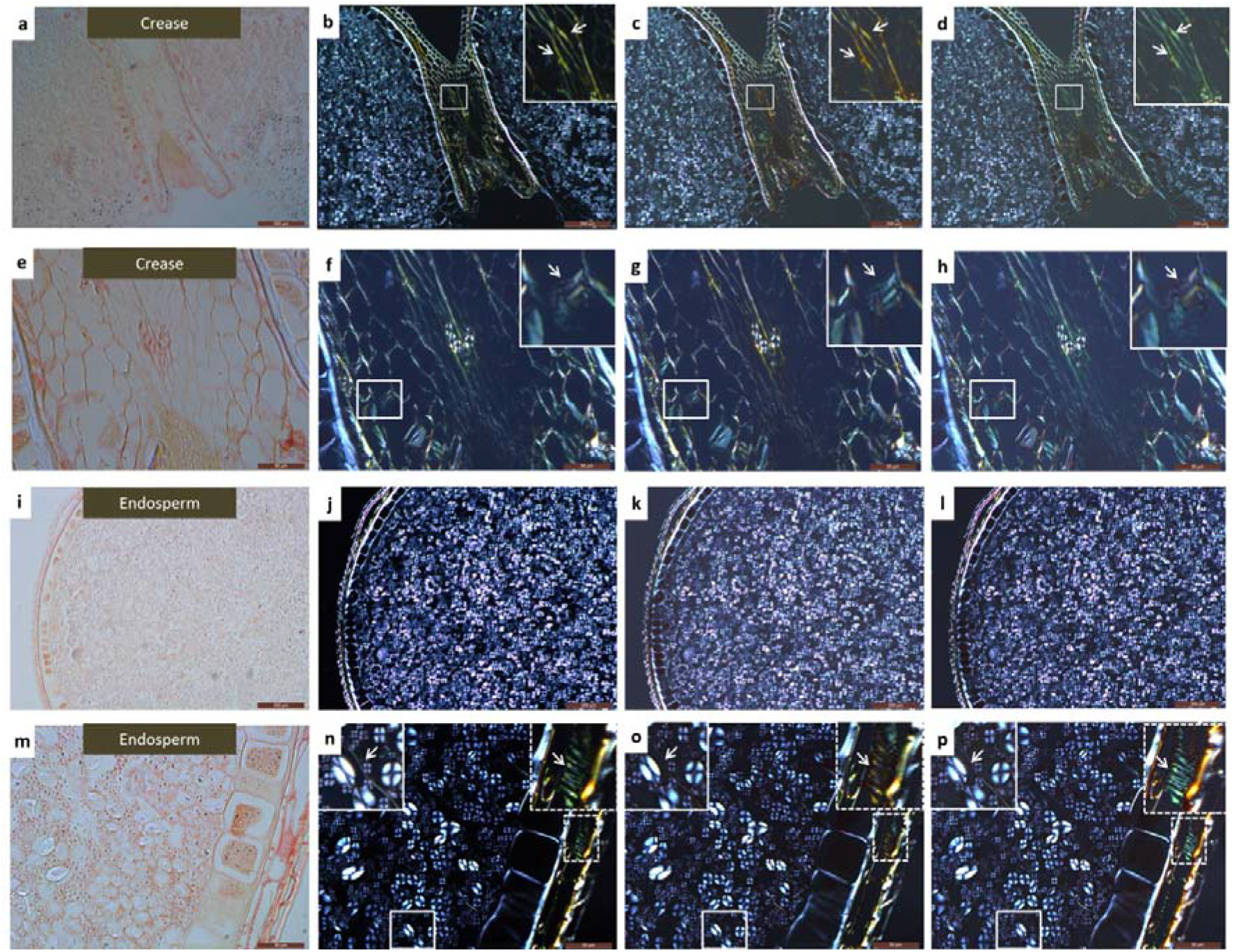
Congo red stained sections of wheat grain crease (a-h) and endosperm (i-p). The crease region shows congophilicity at the cell wall regions of the nucellar projections. The seed coat as well as the endosperm cell walls exhibit congophilicity, suggesting the presence of amyloid-like strctures in these regions. (a-d and i-l represents images obtained with 10x objective, whereas e-h and m-p represents images obtained with 40x objective; the second column represents images with polarized light with angle of polarizer set at 0°; the third column represents images with polarized light with angle of polarizer set at 10° to left; the fourth column represents images with polarized light with angle of polarizer set at 10° to right; Arrows in the insets represent transformative feature of amyloids under polarized light)

#### Acid fuchsin and congo red staining of seed sections

To observe amyloid network further, lentil (*L.culinaris*) grains were sectioned using an ultramicrotome, and the sectioned samples were simultaneously stained with acid fuchsin and congo red. Acid fuchsin stains the proteins to purplish-red under bright field microscope, whereas Congo red intercalates to amyloid-like structures. The images were analysed as to whether Congo red and acid fuchsin are binding to same regions, to demarcate the contributing role of proteins to these amyloid network-like structures. The seed sections of lentils show apple-green birefringence in a scaffold-like form interspersed between the cotyledon cells in the region of the cells walls. As expected, acid fuchsin binds strongly to the protein matrix interspersed in between the starch granules of the cotyledon cells, imparting them with purple-red stain. Congo red used alone binds to the cell wall regions and also shows fluorescence around these areas. However, the scaffold-like networks present around the cell walls, are stained with both acid fuchsin and Congo red, suggesting further the presence of amyloid-like proteinaceous structures in these areas. Lentil cotyledon cell walls are composed of arabinan, arabinogalactan and galacturonic acid similar to other dicots.[40] The cotyledon cell walls of lentils are also composed of 6.5% protein.[41] Similar to monocots such as wheat, lentil or other dicot seeds may also contain protein-polysaccharide associations leading to amyloid-like structure formation in the seeds.

**Figure 4:**
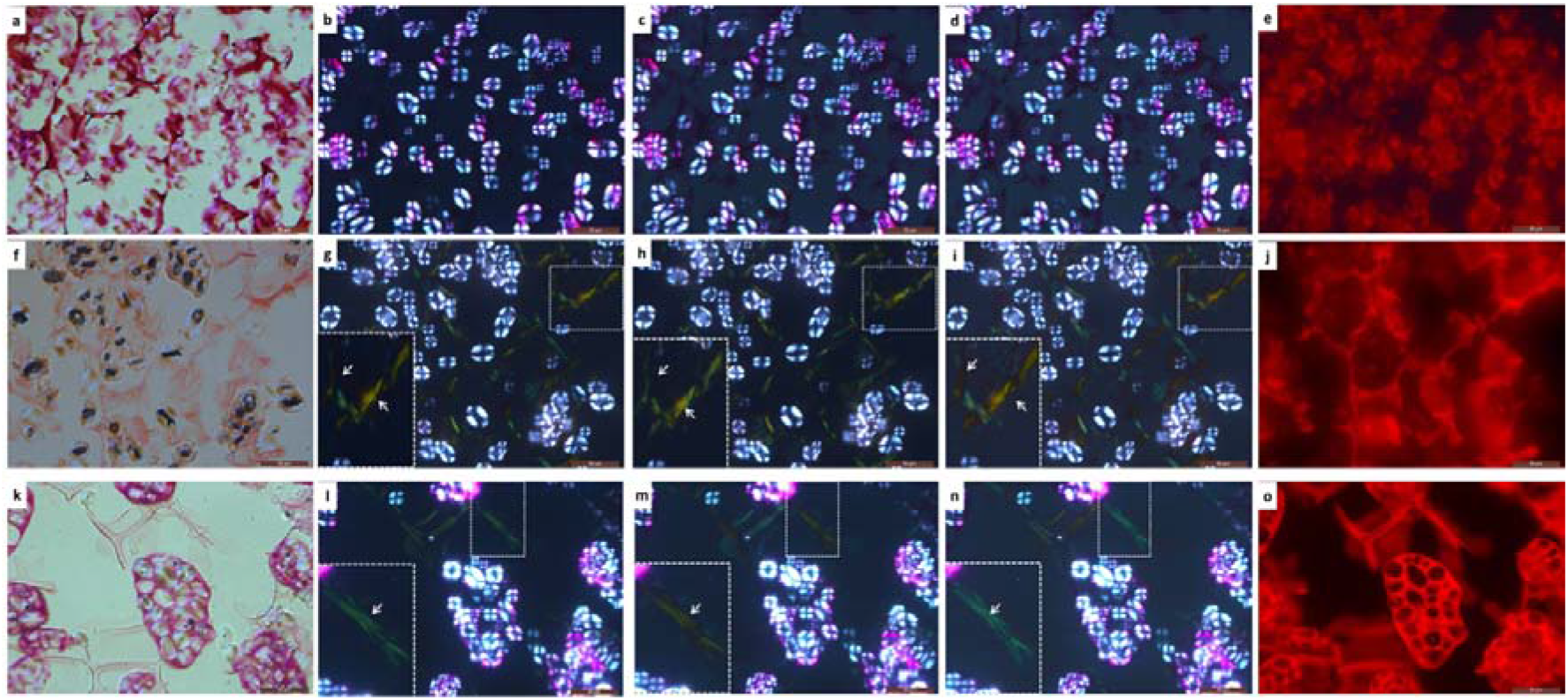
Acid fuchsin (a-e), Congo red (f-j) and acid fuchsin+congo red (k-o) stained sections of lentils. Acid fuchsin stains the cell matrix strongly, but also shows binding to starch granules (a and e) as observed under bright-field and fluorescent microscopy. Congo red stains only the cell wall regions (f and j) and shows apple-green birefringence in these regions (g-i). Simultaneous treatment with both stains however shows stain in the cellular matrix and cell wll regions (k,o) but do not stain starch granules (o) as evident in the fluorescent microscope; apple-green birefringence is still observed in the cell wall regions. (Arrows in the insets represent transformative feature of amyloids under polarized light)

### Comparison between proteinaceous endosperm and starch-rich compartment

To check whether the amyloid network is contributed majorly by the protein-rich or carbohydrate-rich environment, we compared the Congo-red stained sections of potato and wheat seeds. Starch-rich potato sections showed prominent apple-green birefringence at the outermost seed coat regions, signifying that in this peridermal region, cell walls are congophilic. Amyloid birefringence was not observed however in the centre, but there is surplus amounts of starch deposition. The potato cell walls contain arabinose, galactose, galacturonic acid, arabinogalactan, xyloglucan and 1.7% protein.[42] Therefore, comparable carbohydrate and protein content in potato and seed cell walls, signify that the association with type of protein, their sequence or structure might play an important role in congophilicity. The wheat seed sections on the other hand show a congophilic area in the aleurone cell walls, inside aleurone cells and in endosperm cell walls. As discussed before, the aleurone and endosperm cells contain a significant amount of protein in their cell walls as well as contain multiple protein storage vacuoles in their cytoplasm. The characteristic amyloidic signatures in these regions signify the presence of an amyloid network in the cells.

**Figure 5:**
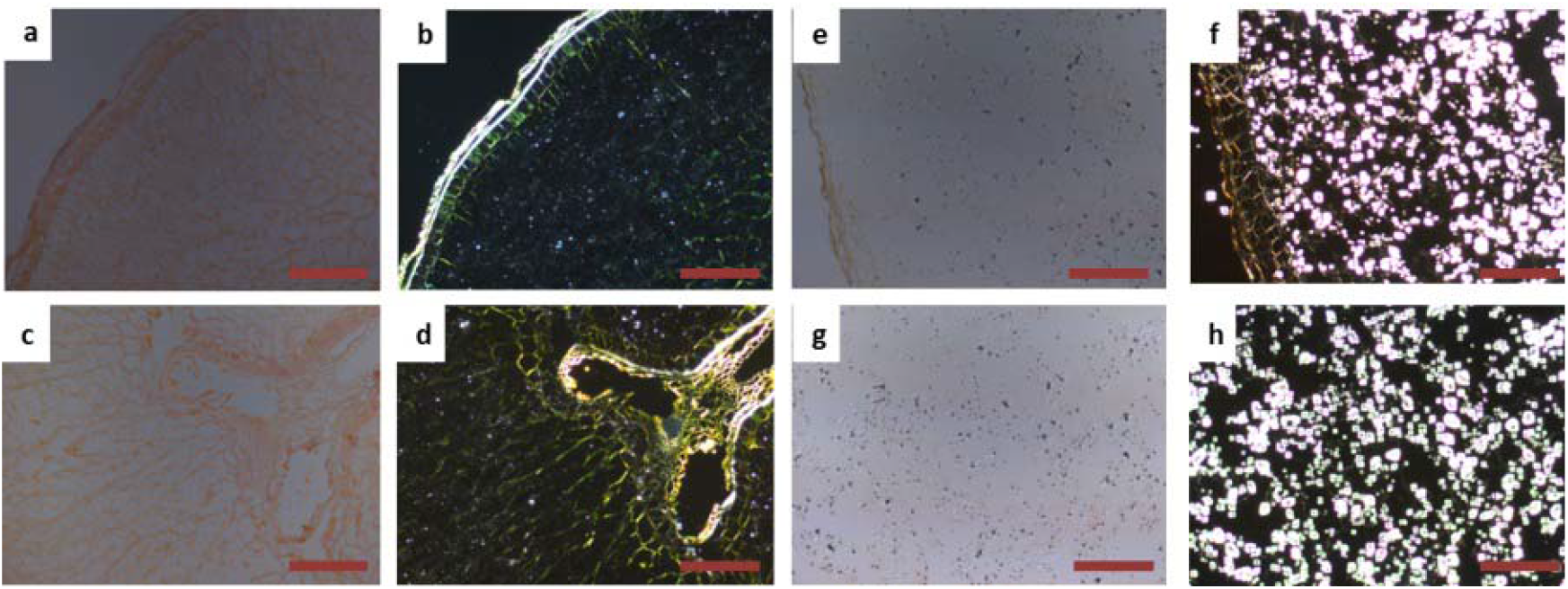
Congo red-stained sections of wheat (a-d) and potato (e-h) shows the presence of Congophilic network prominent in wheat seed sections and dominance of starch granules in potato sections. Scale bar - 200μm

### Congo red and calcofluor staining of isolated cell walls

Since the apple-green birefringence was observed prominently in the cell wall regions, we isolated and characterized the cell walls by staining them with congo red and calcofluor white to demarcate the contributing role of cell walls to the birefringence. Calcoflour white binds to the cellulose and β-glucan molecules present in the cell walls of plants and emits an intense blue fluorescence. In lentil and chickpea seed cell walls, blue fluorescence of calcofluor signified the presence of a large amount of cellulose and glucans. These cell walls also exhibited apple-green birefringence and congo-red binding. On the other hand, wheat showed a milder fluorescence from the calcofluor signifying the lesser amount of cellulose or glucans in this monocot. The amount of birefringence in these areas is also significantly lower than the other seeds. This is possibly because of the compositional differences in the monocot and dicot cell walls.

**Figure 6:**
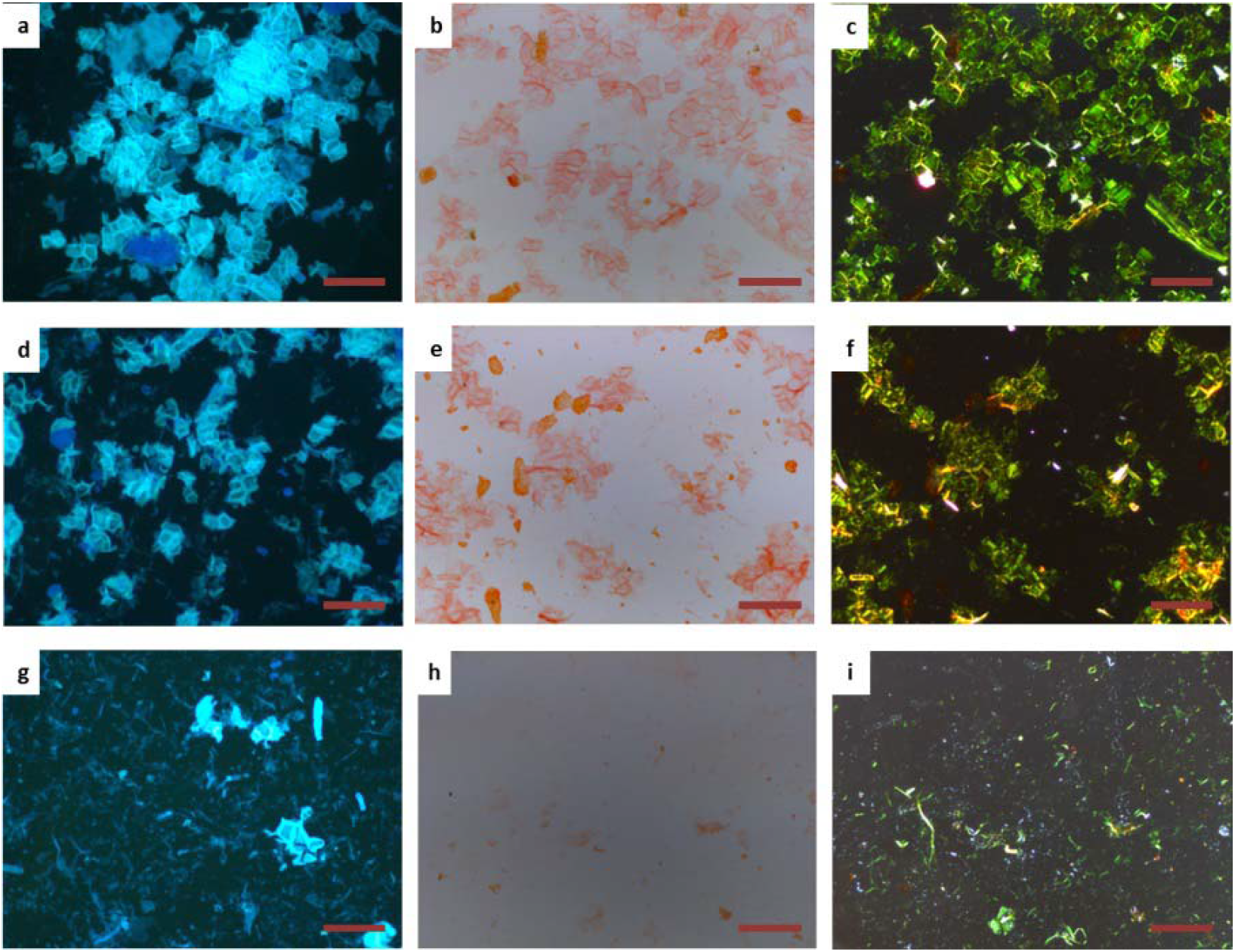
Congo red and calcofluor white stained sections of chickpea (a-c), lentil (d-f), and wheat (g-i). The chickpea and lentil cell walls show apple-green birefringence elucidating the presence of amyloid-like structures even in plant carbohydrates. In wheat seeds, the lack of birefringence in cell walls suggests the compositional difference in the monocot cell walls. Scale bar - 200μm

#### Conclusion

Although amyloids are known to be associated with carbohydrates and other biomolecules, not much is known about these associations in the plant tissues. We hypothesized that the plant seeds might exhibit such heterogeneous amyloid networks due to the presence of protein-rich storage granules and their close association with starch molecules in seed endosperm or cotyledon cells. The protein bodies are rich in glutamine-rich proteins, which might impart aggragation propensity to these molecules. In this paper, we have shown that plant seeds have a mesh-like network that is congophilic. Therefore, we propose that plants have an amyloid-like network composed of proteins, carbohydrates, and other biomolecules associated together, forming a complex interactome. The roles of such structures might involve structural as well as functional and needs further investigation. Further work in this direction such as exact compositional analysis of this network is in progress.

## Materials and methods

### Materials used

Mature seeds were collected from the local market. Glutaraldehyde, ethanol, and xylene were obtained from Thermo Scientific. Paraformaldehyde, Congo red (CR) thioflavin-T (ThT), acid fuchsin, and calcofluor white were obtained from Sigma Aldrich. Paraffin wax for embedding was obtained from Leica. Microscopic slides were from HiMedia. The sections were prepared using Leica Microtome. The tissue sections were visualized using a Leica DM 2500 fluorescent microscope equipped with cross-polarizers.

### Seed sectioning

A fixation solution was prepared to contain 2.5% glutaraldehyde and 3% paraformaldehyde in 0.1 M phosphate buffer. The pH of the fixative solution was maintained at 7.2. The seeds were incubated in this solution for 2 hours at room temperature (24-26°C) and were then transferred to 4°C for overnight fixation. The seeds were then washed three times with phosphate buffer solution. Dehydration of the seeds was then performed with successive gradients of ethanol (30%, 50%, 70%, 90%) for 30 minutes each. This was followed by dehydration at 100% ethanol twice for 1 hour each. The samples were then permeated with xylene and finally with paraffin. Paraffin embedding was continued overnight at 60°C. The samples were then cast into blocks and stored at 4°C until further processing. For sectioning, the embedded blocks were attached to the Leica microtome trimmer. Extra wax was first cut by a razor blade, and the microtome was then used to obtain sections of 8 μm thickness. The sections were carefully transferred in water, followed by a glass slide. The slides were then heated on a stage for 30 minutes at 50°C.

### Staining techniques

Congo red staining of the samples was done using saturated congo red solution filtered through 45 μm filter. For thioflavin-T staining, 20 μM aqueous ThT solution was prepared. The slides were stained with the respective dye, air-dried, and repeatedly washed with distilled water. Birefringence and Congo red binding were observed using bright-field microscopy and polarized light.[43] ThT-stain was observed using the 660 nm red filter of a fluorescence microscope. For acid fuchsin stain preparation, 3.5 g dye and 250 ml glacial acetic acid are added to 750 ml distilled water. The stained sections are then observed under bright-field microscopy. A stock solution of calcofluor white stain was prepared by adding 35 mg of the dye to 7 ml distilled water and 100 μl 10 N NaOH. The volume was adjusted to 10 ml with distilled water. From this stock, a 10% vol/vol solution was made for further experiments. The sections were visualized under the UV filter of a fluorescent microscope.

## References

[1] A. Aguzzi, T. O’Connor, Protein aggregation diseases: pathogenicity and therapeutic perspectives, Nature Reviews Drug Discovery 9(3) (2010) 237–248.

[2] J.D. Sipe, M.D. Benson, J.N. Buxbaum, S.I. Ikeda, G. Merlini, M.J. Saraiva, P. Westermark, Amyloid fibril proteins and amyloidosis: chemical identification and clinical classification International Society of Amyloidosis 2016 Nomenclature Guidelines, Amyloid : the international journal of experimental and clinical investigation : the official journal of the International Society of Amyloidosis 23(4) (2016) 209–213.

[3] A.J. Howie, “Green (or apple-green) birefringence” of Congo red-stained amyloid, Amyloid : the international journal of experimental and clinical investigation : the official journal of the International Society of Amyloidosis 22(3) (2015) 205–6.

[4] P. Westermark, Aspects on human amyloid forms and their fibril polypeptides, The FEBS Journal 272(23) (2005) 5942–5949.

[5] M. Ankarcrona, B. Winblad, C. Monteiro, C. Fearns, E.T. Powers, J. Johansson, G.T. Westermark, J. Presto, B.G. Ericzon, J.W. Kelly, Current and future treatment of amyloid diseases, J Intern Med 280(2) (2016) 177–202.

[6] M.R. Chapman, L.S. Robinson, J.S. Pinkner, R. Roth, J. Heuser, M. Hammar, S. Normark, S.J. Hultgren, Role of Escherichia coli curli operons in directing amyloid fiber formation, Science (New York, N.Y.) 295(5556) (2002) 851–5.

[7] C.B. Ramsook, C. Tan, M.C. Garcia, R. Fung, G. Soybelman, R. Henry, A. Litewka, S. O’Meally, H.N. Otoo, R.A. Khalaf, A.M. Dranginis, N.K. Gaur, S.A. Klotz, J.M. Rauceo, C.K. Jue, P.N. Lipke, Yeast Cell Adhesion Molecules Have Functional Amyloid-Forming Sequences, Eukaryotic Cell 9(3) (2010) 393–404.

[8] D.M. Fowler, A.V. Koulov, C. Alory-Jost, M.S. Marks, W.E. Balch, J.W. Kelly, Functional amyloid formation within mammalian tissue, PLoS Biol 4(1) (2006) e6.

[9] M.P. Jackson, E.W. Hewitt, Why are Functional Amyloids Non-Toxic in Humans?, Biomolecules 7(4) (2017) 71.

[10] R.A. Kyle, Amyloidosis: a convoluted story, British Journal of Haematology 114(3) (2001) 529–538.

[11] R. Gallardo, N.A. Ranson, S.E. Radford, Amyloid structures: much more than just a cross-β fold, Current Opinion in Structural Biology 60 (2020) 7–16.

[12] K.L. Stewart, E. Hughes, E.A. Yates, G.R. Akien, T.-Y. Huang, M.A. Lima, T.R. Rudd, M. Guerrini, S.-C. Hung, S.E. Radford, D.A. Middleton, Atomic Details of the Interactions of Glycosaminoglycans with Amyloid-β Fibrils, Journal of the American Chemical Society 138(27) (2016) 8328–8331.

[13] X. Zhan, B. Stamova, F.R. Sharp, Lipopolysaccharide Associates with Amyloid Plaques, Neurons and Oligodendrocytes in Alzheimer’s Disease Brain: A Review, Frontiers in aging neuroscience 10 (2018) 42.

[14] R.A. Dluhy, S. Shanmukh, J.B. Leapard, P. Krüger, J.E. Baatz, Deacylated pulmonary surfactant protein SP-C transforms from alpha-helical to amyloid fibril structure via a pH-dependent mechanism: an infrared structural investigation, Biophys J 85(4) (2003) 2417–29.

[15] M. Stefani, Biochemical and biophysical features of both oligomer/fibril and cell membrane in amyloid cytotoxicity, The FEBS Journal 277(22) (2010) 4602–4613.

[16] T. Bitter, H. Muir, Mucopolysaccharides of whole human spleens in generalized amyloidosis, The Journal of clinical investigation 45(6) (1966) 963–75.

[17] D.J. Martin, M. Ramirez-Alvarado, Glycosaminoglycans promote fibril formation by amyloidogenic immunoglobulin light chains through a transient interaction, Biophys Chem 158(1) (2011) 81–9.

[18] J.J. Aguilera, F. Zhang, J.M. Beaudet, R.J. Linhardt, W. Colón, Divergent effect of glycosaminoglycans on the in vitro aggregation of serum amyloid A, Biochimie 104 (2014) 70–80.

[19] D.O. Serra, A.M. Richter, R. Hengge, Cellulose as an architectural element in spatially structured Escherichia coli biofilms, Journal of bacteriology 195(24) (2013) 5540–54.

[20] D. Laor, D. Sade, S. Shaham-Niv, D. Zaguri, M. Gartner, V. Basavalingappa, A. Raveh, E. Pichinuk, H. Engel, K. Iwasaki, T. Yamamoto, H. Noothalapati, E. Gazit, Fibril formation and therapeutic targeting of amyloid-like structures in a yeast model of adenine accumulation, Nature Communications 10(1) (2019) 62.

[21] V. Singh, R.K. Rai, A. Arora, N. Sinha, A.K. Thakur, Therapeutic implication of L-phenylalanine aggregation mechanism and its modulation by D-phenylalanine in phenylketonuria, Scientific Reports 4(1) (2014) 3875.

[22] P. Wood, R. Fulcher, Interaction of some dyes with cereal β-Glucans, Cereal chemistry Nov/Dec 55 (1978) 952–966.

[23] Verbelen, Kerstens, Polarization confocal microscopy and Congo Red fluorescence: a simple and rapid method to determine the mean cellulose fibril orientation in plants, Journal of Microscopy 198(2) (2000) 101–107.

[24] P.R. Shewry, N.G. Halford, Cereal seed storage proteins: structures, properties and role in grain utilization, Journal of experimental botany 53(370) (2002) 947–958.

[25] A.A.M. Andersson, R. Andersson, K. Autio, P. Åman, Chemical Composition and Microstructure of Two Naked Waxy Barleys, Journal of Cereal Science 30(2) (1999) 183–191.

[26] A.V. Balakireva, A.A. Zamyatnin, Properties of Gluten Intolerance: Gluten Structure, Evolution, Pathogenicity and Detoxification Capabilities, Nutrients 8(10) (2016) 644.

[27] H.C. Fan, L.I. Ho, C.S. Chi, S.J. Chen, G.S. Peng, T.M. Chan, S.Z. Lin, H.J. Harn, Polyglutamine (PolyQ) diseases: genetics to treatments, Cell Transplant 23(4-5) (2014) 441–58.

[28] M. Monge-Morera, M.A. Lambrecht, L.J. Deleu, N.N. Louros, F. Rousseau, J. Schymkowitz, J.A. Delcour, Heating Wheat Gluten Promotes the Formation of Amyloid-like Fibrils, ACS Omega (2021).

[29] K.S. Antonets, A.A. Nizhnikov, Amyloids and prions in plants: Facts and perspectives, Prion 11(5) (2017) 300–312.

[30] P.J. Langston-Unkefer, W. Gade, A Seed Storage Protein with Possible Self-Affinity through Lectin-Like Binding, Plant Physiology 74(3) (1984) 675–680.

[31] N.J. Atkin, S.L. Cheng, R.M. Abeysekera, A.W. Robards, Localisation of Amylose and Amylopectin in Starch Granules Using Enzyme-Gold Labelling, Starch - Stärke 51(5) (1999) 163–172.

[32] L.H. Thomas, V.T. Forsyth, A. Šturcová, C.J. Kennedy, R.P. May, C.M. Altaner, D.C. Apperley, T.J. Wess, M.C. Jarvis, Structure of Cellulose Microfibrils in Primary Cell Walls from Collenchyma, Plant Physiology 161(1) (2013) 465.

[33] M. Dahesh, A. Banc, A. Duri, M.H. Morel, L. Ramos, Polymeric assembly of gluten proteins in an aqueous ethanol solvent, The journal of physical chemistry. B 118(38) (2014) 11065–76.

[34] K.J.A. Jansens, M.A. Lambrecht, I. Rombouts, M. Monge Morera, K. Brijs, F. Rousseau, J. Schymkowitz, J.A. Delcour, Conditions Governing Food Protein Amyloid Fibril Formation—Part I: Egg and Cereal Proteins, Comprehensive Reviews in Food Science and Food Safety 18(4) (2019) 1256–1276.

[35] K.S. Antonets, M.V. Belousov, A.I. Sulatskaya, M.E. Belousova, A.O. Kosolapova, M.I. Sulatsky, E.A. Andreeva, P.A. Zykin, Y.V. Malovichko, O.Y. Shtark, A.N. Lykholay, K.V. Volkov, I.M. Kuznetsova, K.K. Turoverov, E.Y. Kochetkova, A.G. Bobylev, K.S. Usachev, O.N. Demidov, I.A. Tikhonovich, A.A. Nizhnikov, Accumulation of storage proteins in plant seeds is mediated by amyloid formation, PLoS Biol 18(7) (2020) e3000564.

[36] P.R. Shewry, 7 - Improving the content and composition of dietary fibre in wheat, in: J.A. Delcour, K. Poutanen (Eds.), Fibre-Rich and Wholegrain Foods, Woodhead Publishing2013, pp. 153–169.

[37] D.I. Rhodes, B.A. Stone, Proteins in Walls of Wheat Aleurone Cells, Journal of Cereal Science 36(1) (2002) 83–101.

[38] R. Palmer, V. Cornuault, S.E. Marcus, J.P. Knox, P.R. Shewry, P. Tosi, Comparative in situ analyses of cell wall matrix polysaccharide dynamics in developing rice and wheat grain, Planta 241(3) (2015) 669–685.

[39] A.M. Showalter, Arabinogalactan-proteins: structure, expression and function, Cellular and Molecular Life Sciences CMLS 58(10) (2001) 1399–1417.

[40] T.M. Shiga, F.M. Lajolo, T.M.C.C. Filisetti, Cell wall polysaccharides of common beans (Phaseolus vulgaris L.), Food Science and Technology 23 (2003) 141–148.

[41] R.S. Bhatty, Cooking quality of lentils: the role of structure and composition of cell walls, Journal of Agricultural and Food Chemistry 38(2) (1990) 376–383.

[42] M.C. Jarvis, M.A. Hall, D.R. Threlfall, J. Friend, The polysaccharide structure of potato cell walls: Chemical fractionation, Planta 152(2) (1981) 93–100.

[43] L. Zhao, T. Pan, C. Cai, J. Wang, C. Wei, Application of whole sections of mature cereal seeds to visualize the morphology of endosperm cell and starch and the distribution of storage protein, Journal of Cereal Science 71 (2016) 19–27.

